# Independent control of neurogenesis and dorsoventral patterning by NKX2-2

**DOI:** 10.1101/2024.10.13.618113

**Authors:** Sumin Jang, Elena Abarinov, Julie Dobkin, Hynek Wichterle

## Abstract

Human neurogenesis is disproportionately protracted, lasting >10 times longer than in mouse, allowing neural progenitors to undergo more rounds of self-renewing cell divisions and generate larger neuronal populations. In the human spinal cord, expansion of the motor neuron lineage is achieved through a newly evolved progenitor domain called vpMN (ventral motor neuron progenitor) that uniquely extends and expands motor neurogenesis. This behavior of vpMNs is controlled by transcription factor NKX2-2, which in vpMNs is co-expressed with classical motor neuron progenitor (pMN) marker OLIG2. In this study, we sought to determine the molecular basis of NKX2-2-mediated extension and expansion of motor neurogenesis. We found that NKX2-2 represses proneural gene *NEUROG2* by two distinct, Notch-independent mechanisms that are respectively apparent in rodent and human spinal progenitors: in rodents (and chick), NKX2-2 represses *Olig2* and the motor neuron lineage through its tinman domain, leading to loss of *Neurog2* expression. In human vpMNs, however, NKX2-2 represses *NEUROG2* but not *OLIG2*, thereby allowing motor neurogenesis to proceed, albeit in a delayed and protracted manner. Interestingly, we found that ectopic expression of tinman-mutant *Nkx2-2* in mouse pMNs phenocopies human vpMNs, repressing *Neurog2* but not *Olig2*, and leading to delayed and protracted motor neurogenesis. Our studies identify a Notch- and tinman-independent mode of *Nkx2-2*-mediated *Neurog2* repression that is observed in human spinal progenitors, but is normally masked in rodents and chicks due to *Nkx2-2*’s tinman-dependent repression of *Olig2*.

## INTRODUCTION

The central nervous system (CNS) is disproportionately expanded in humans compared to other mammals like rodents and non-human primates, underscoring the increased range and complexity of our cognitive and behavioral capacity^1^. The size of the nervous system is directly impacted by the mitotic behavior of neural progenitors during development^2^. At each cell cycle during neurogenesis, neural progenitors either choose to re-enter the cell cycle and self-renew, or exit the cell cycle and acquire a neuronal fate. The balance between these two alternative decisions directly affects how many neurons are generated during the neurogenic period.

Several signaling molecules have been shown to change the mitotic behavior of neural progenitors and increase their tendency to re-enter the cell cycle and self-renew^3–7^. One such signal that operates in the vast majority of neural lineages is the Notch signaling pathway^8^. Activation of Notch leads to expression of downstream effector genes belonging to the Hes/Hey family, which represses a group of proneural genes that include Neurogenins (Neurog)^9^. Neurogenins have been shown to be necessary for neuronal fate commitment of progenitors across a wide range of lineages within the CNS^10^. Indeed, pharmacological inactivation of Notch signaling (using gamma secretase inhibitor DAPT, for example) sharply increases Neurog expression and rapidly prompts neural progenitors to exit the cell cycle, while stable activation of Notch (e.g. by expression of Notch Intracellular Domain, NICD) inhibits neurogenesis^11^. In addition to repressing Neurog, Hes genes also represses their own expression, leading to the off-phase oscillatory expression of Hes and Neurog genes in neural progenitors, known as the Hes cycle^12^ **(Figure 1B)**. Depending on the phase of the Hes cycle, progenitors are biased either toward remaining mitotic (Hes-high, Neurog-low), or undergoing neurogenesis (Hes-low, Neurog-high). Although the Notch signaling pathway is highly conserved across animals, it has also undergone evolutionary modifications that impact the behavior of neural progenitors, contributing to the marked expansion of the CNS in primates. Until recently, the vast majority of studies on human neural expansion focused on the neocortex^13,14^, where, among others, expression of human-specific Notch receptor paralog *NOTCH2NL* was shown to bias human progenitors to remain in cell cycle for longer, thus expanding neurogenic output^6,7^.

**Figure 1:**
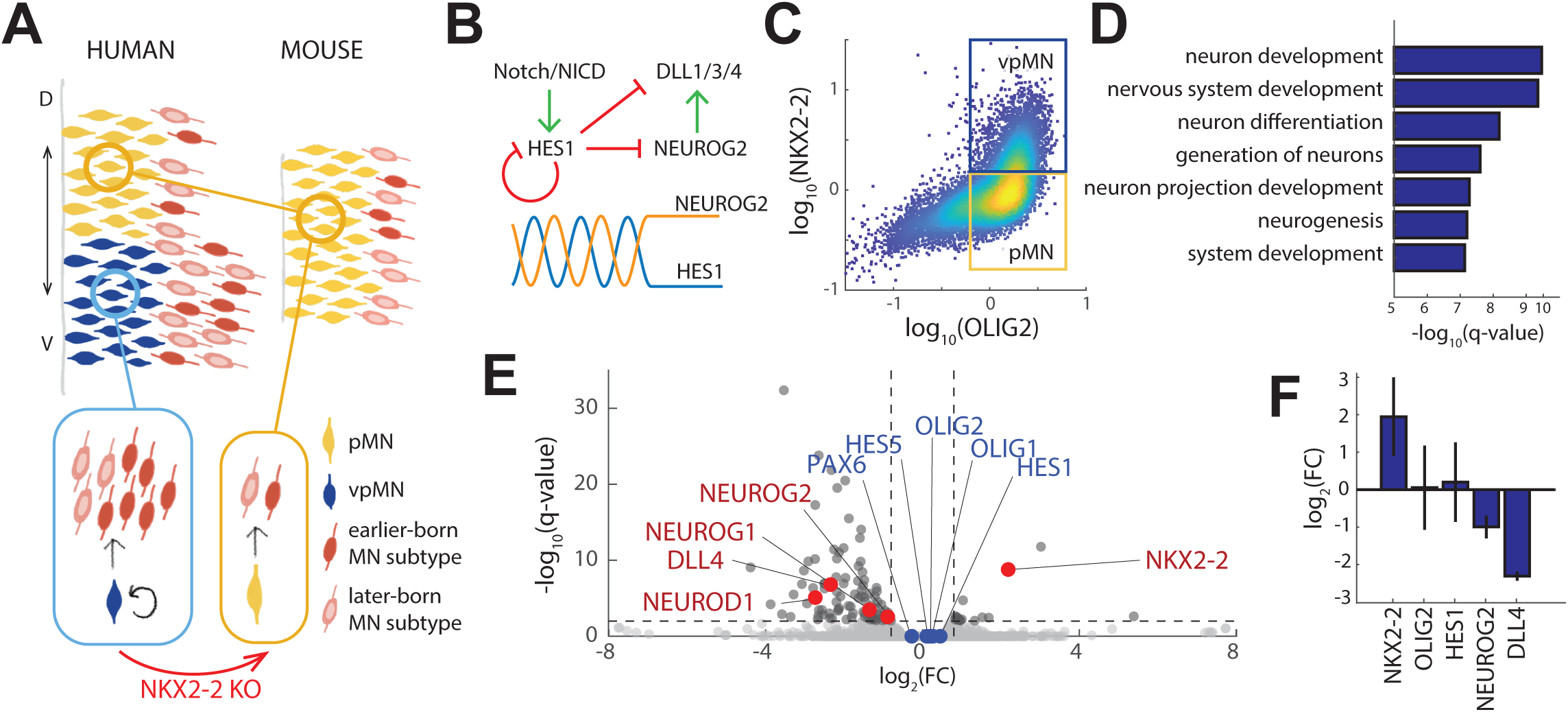
NKX2-2 represses NEUROG2 in vpMNs in a HES-independent manner. **(A)** Schematic of human and mouse motor neurogenesis. Human motor neurogenesis involves a second, newly-evolved vpMN domain that uniquely expands motor neuron production, which is functionally dependent on NKX2-2. **(B)** Notch signaling activates Hes, which represses proneural genes Neurog in an oscillatory manner, giving rise to the Hes cycle. **(C)** Flow cytometry density scatter plot of day 11 human cultures immunostained for NKX2-2 and OLIG2. vpMN and pMNs were purified for subsequent RNA-seq. **(D)** GO-term enrichment analysis results for genes significantly (q-value < 0.01) downregulated in vpMNs compared to pMNs. **(E)** vpMNs display decreased expression of *NEUROG2* and downstream target genes, while *OLIG2*, *PAX6*, and *HES1/5* remain unchanged. **(F)** *NKX2-2* knockout leads to increased *NEUROG2* but does not affect *OLIG2* or *HES* expression in vpMNs.

In the developing spinal cord, evolutionary expansion of motor neurons relies on a newly evolved ventral motor neuron progenitor (vpMN) domain that is present in humans but absent in rodents and chicks^15^. These “human-specific” vpMNs uniquely undergo protracted neurogenesis, giving rise to ∼10 motor neurons per progenitor, while the conserved motor neuron progenitors (pMNs) give rise on average to ∼2 motor neurons in both humans and mice **(Figure 1A)**. vpMNs are characterized by co-expression of *NKX2-2* and *OLIG2*, two transcription factors that in rodents and chicks are expressed in adjacent but non-overlapping progenitor domains: *Nkx2-2* is expressed in the ventral-most p3 progenitor domain, while *Olig2* is expressed in the pMN domain, which lies immediately dorsal to p3^16,17^. Importantly, the extended neurogenic behavior of vpMNs is dependent on *NKX2-2* expression, as knockout of *NKX2-2* converts these progenitors to pMN-like cells, giving rise only to two motor neurons^15^ **(Figure 1A)**. Interestingly, while vpMNs exhibit increased self-renewal, they do not express higher level of *NOTCH2NL* compared to pMNs. This raises several important questions: What are the mechanisms by which *NKX2-*2 alters the mitotic behavior of vpMNs? And can forced expression of *Nkx2-2* in mouse motor neuron progenitors humanize their behavior, leading to prolonged neurogenesis and increased motor neuron production?

We hypothesized that NKX2-2 might exert a complex effect on neural progenitors, mediated by different conserved functional domains found in this transcription factor. In order to investigate this possibility, we took advantage of inducible mouse embryonic stem cells (ESCs) that were engineered to express either WT *Nkx2-2* or mutant versions of the transcription factor, in which either (a) the Groucho co-repressor recruiting tinman (TN) domain^18^, or (b) the conserved NK2-specific domain (SD) that mediates protein-protein interactions and DNA-binding of Nkx2-2^19,20^, is disrupted. Analysis of gene expression changes in mouse motor neuron progenitors forcibly expressing these transcription factors confirmed that WT Nkx2-2 effectively represses a number of patterning transcription factors, including *Olig2*, *Pax6* and *Irx3*, which specify motor neuron progenitor (pMN) and/or interneuron progenitor (p2) domains that lie dorsal to the p3 domain^21^. As previously reported, this repression was fully dependent on the presence of the TN domain^22^, but not on the SD. Interestingly, while *Nkx2-2* carrying mutations in the TN domain failed to repress patterning genes like *Olig2*, it still exerted repressive activity toward *Neurog2*, a proneural gene that drives differentiation of OLIG2^+^ progenitors into motor neurons. In turn, motor neuron progenitors remained mitotic for longer, extending their neurogenic period and mimicking the behavior of human-specific vpMNs. Thus, our experiments suggest that NKX2-2 has two biochemically distinct functions in neural progenitors. Through its TN domain it represses determinants of motor neuron progenitors and intermediate spinal progenitors, effectively controlling dorsoventral spinal patterning. In the absence of the TN domain, its patterning activity is abolished, but NKX2-2 retains its ability to suppress neurogenesis through a TN-independent mechanism. We speculate that in human spinal progenitors, NKX2-2 is biased towards this TN-independent repressive activity, allowing co-expression of *NKX2-2* with *OLIG2* while extending neurogenesis through *NEUROG2* repression, leading to increased motor neuron production.

## RESULTS

### NKX2-2 represses *NEUROG2* in a HES-independent manner

To investigate the mechanisms downstream of NKX2-2 that lead to delayed and protracted neurogenesis in vpMNs, we performed differential gene expression analysis on vpMNs (NKX2-2^+^/OLIG2^+^) and pMNs (NKX2-2^−^/OLIG2^+^) purified from in vitro differentiated induced pluripotent stem cells (iPSCs) by fluorescent activated sorting of cells immunostained for NKX2-2 and OLIG2 **(Figure 1C)**. We identified 97 genes that were significantly (q-value < 0.01) down-regulated in vpMNs compared to pMNs. In contrast, only 18 genes were upregulated, suggesting that NKX2-2 predominantly operates as a transcriptional repressor, as previous studies have shown^22^.

To gain insight on which pathways NKX2-2 might be acting through to extend motor neurogenesis, we first performed GO-term enrichment analysis on genes that are significantly down-regulated in vpMNs. We found that terms relating to neurogenesis were highly represented in the most enriched pathways **(Figure 1D)**. Among these genes significantly downregulated in vpMNs, we found *NEUROG2*, its immediate downstream targets *DLL4* and *NEUROD1* **(Figure 1E)**, as well as postmitotic motor neuron genes (*MNX1*, *ONECUT1*, *LHX4*) and pan-neuronal genes (*ELAVL3*, *NOVA1*, *STMN2*) **(Table 1)** ^10,23^. Although *NEUROG2* and its downstream targets were significantly downregulated in vpMNs, *HES1/5* expression remained unchanged, indicating similar Notch activity between the two progenitor populations. We also found that *OLIG2* expression remained unchanged between vpMNs and pMNs, in accordance with the stable co-expression of NKX2-2 and OLIG2 observed previously in human motor neurogenesis^15,24,25^. Surprisingly, expression of *PAX6*, which has been shown to be directly repressed by NKX2-2 in chick and mouse^16^, was also unchanged between vpMN and pMN.

We then asked whether NKX2-2 was functionally involved in repressing *NEUROG2* expression. To address this, we knocked out *NKX2-2* in human iPSCs, and saw that expression of *NEUROG2* and *DLL4* indeed increased, while *HES1/5* and *OLIG2* expression levels were unaffected in motor neuron progenitors **(Figure 1F)**. Together, these results suggest that NKX2-2 represses *NEUROG2* in vpMNs independently of dorsoventral patterning genes *OLIG2* and *PAX6*, or the Notch signaling pathway.

### NKX2-2 represses *Neurog2* independently of its tinman domain in upper cervical spinal progenitors

To test whether NKX2-2 is sufficient to repress *Neurog2* and recapitulate the delayed and protracted neurogenic timeline of vpMNs in mouse progenitors, we generated a mouse ESC line harboring a doxycycline (DOX)-inducible *Nkx2-2* transgene in the *HPRT* locus **(Figure 2A)**. To test the role of *Nkx2-2*’s conserved functional domains, we also generated mouse ESC lines carrying DOX-inducible *Nkx2-2* with a mutated TN (TN_mut_) domain, and utilized NK2-specific domain-mutant (SD_mut_) mouse ESC lines generated from previous studies^20^ (hereon referred to as iNkx2-2^WT^, iNkx2-2^TNmut^ or iNkx2-2^SDmut^). Upon treatment of these cells with DOX, we saw rapid and uniform induction of NKX2-2 protein in all three cell lines **(Figure 2B)**.

**Figure 2:**
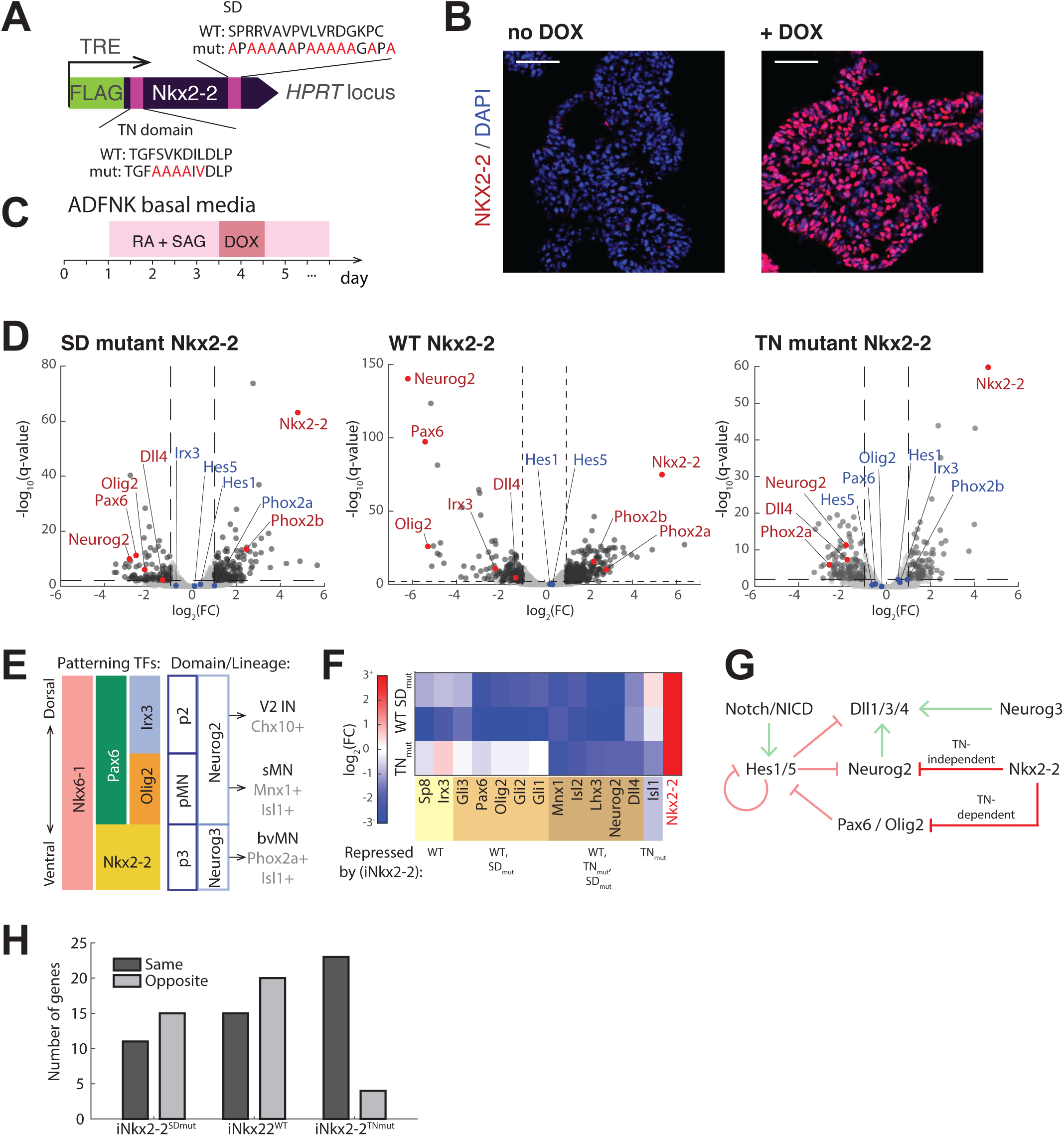
NKX2-2 represses *Neurog2* independently of its tinman domain in upper cervical spinal progenitors. **(A)** Schematic of DOX-inducible constructs containing WT, TN-mutant or SD-mutant Nkx2-2**. (B)** DOX treatment leads to uniform ectopic expression of NKX2-2 protein. **(C)** Summary of culture conditions that produce rostral cervical motor neurons. **(D)** Differential gene expression analysis between non-DOX-treated and iNkx2-2^SDmut^ (left), iNkx2-2^WT^ (middle) or iNkx2-2^TNmut^ cells on day 4. **(E)** Schematic of ventral progenitor domains in the rostral cervical spinal cord, along with patterning genes, proneural genes and neuronal fate for each lineage. (IN = interneuron, sMN = somatic motor neuron, bvMN = branchiovisceral motor neuron) **(F)** Expression level changes of patterning genes, proneural and immediate downstream target genes, and somatic/branchiovisceral motor neuron genes between non-DOX treated and iNkx2-2^SDmut^, iNkx2-2^WT^, or iNkx2-2^TNmut^ cells on day 4. pMN and p2 lineage genes are repressed, while p3 genes are unregulated by WT and SD-mutant *Nkx2-2*, whereas proneural and somatic motor neuron genes are repressed by all three forms of *Nkx2-2*. *Isl1*, which is expressed in both somatic and branchiovisceral motor neurons, is repressed only by iNkx2-2^TNmut^. **(G)** *Nkx2-2* represses *Neurog2* through two distinct, Notch-independent mechanisms; by repressing *Olig2* through its TN domain, as well as independently of its TN domain and *Olig2*. **(H)** Classification of genes that show significant expression change between pMNs versus vpMNs in human, and non-DOX-treated control versus iNkx2-2^SDmut^, iNkx2-2^WT^, or iNkx2-2^TNmut^ in mouse. iNkx2-2^TNmut^ shows the largest number of genes that show the same direction of change between human and mouse, and the smallest number of genes showing the opposite direction of change, suggesting that iNkx2-2^TNmut^ most similarly recapitulates the molecular differences between vpMNs and pMNs.

We then differentiated the inducible ES cells into the motor neuron lineage using inductive signals retinoic acid (RA) and smoothened agonist (SAG) **(Figure 2C)**. Under these conditions, progenitors acquire a rostral cervical (HOXA5^+^) identity and differentiate into mostly (>90%) somatic (MNX1^+^/ISL1^+^) and some (<10%) branchiovisceral (PHOX2A/B^+^/ISL1^+^) motor neurons^26^, which derive from OLIG2^+^ pMN and NKX2-2^+^ p3 progenitors, respectively^27^ **(Figure 2E)**. Expression of *Nkx2-2* was induced with DOX on day 3.5, 12 hours prior to OLIG2^+^ pMN specification, and cells were harvested 24 hours later for RNA-seq.

Compared to the non-DOX-treated control, iNkx2-2^WT^ and iNkx2-2^SDmut^ showed strong downregulation of somatic motor neuron progenitor (*Olig2* and *Pax6*) and somatic motor neuron (*Mnx1*, *Lhx3/4*) markers, and a concomitant increase of branchiovisceral lineage markers *Phox2a/b* **(Figure 2D, E)**. Meanwhile, these genes were unaffected in iNkx2-2^TNmut^ **(Figure 2D)**, confirming that NKX2-2-mediated repression of pMN and p2 lineage genes requires the TN domain^22^. Ectopic expression of *Nkx2-2* therefore ventralized the gene expression program toward the p3 lineage, while suppressing more dorsal pMN and p2 lineages, in a TN-dependent manner.

Although we initially expected that NKX2-2’s repressive function depends on its TN domain^20^, we observed a set of large set of genes that significantly (q-value < 0.05) changed their expression following induction of TN-mutant *Nkx2-2* (290 upregulated and 317 downregulated). Furthermore, we found that iNkx2-2^WT^ and iNkx2-2^TNmut^ had distinct effects on gene expression (R^2^ = 0.2840), with >15% of significantly affected genes changing expression levels in opposite directions in one versus the other. In contrast, the effects of iNkx2-2^WT^ and iNkx2-2^SDmut^ were highly correlated (R^2^ = 0.7253), with <1% of significantly affected genes changing in opposite directions. Interestingly, Neurog2 was among genes significantly downregulated by iNkx2-2^TNmut^ (as well as by iNkx2-2^WT^), along with downstream targets such as Dll4 and somatic motor neuron (sMN) genes Mnx1 and Lhx3 **(Figure 2F)**. However, the repression of somatic motor neurogenesis arises from two distinct mechanisms in iNkx2-2^WT^ and iNkx2-2^TNmut^. In iNkx2-2^WT^, repression of sMN genes were accompanied by a concomitant increase in branchiovisceral motor neuron (bvMN) genes *Phox2a/b*, indicating that repression of somatic motor neurogenesis arises mainly through ventralization of the lineage identity toward p3. In contrast, iNkx2-2^TNmut^ repressed *Neurog2* and somatic motor neurogenesis without altering lineage identity. Indeed, *Isl1*, which is expressed in both somatic and branchiovisceral motor neurons, is repressed in iNkx2-2^TNmut^ but unaffected in iNkx2-2^WT^ or iNkx2-2^SDmut^ **(Figure 2D, F)**. Together, these results reveal the dual functions of NKX2-2 in spinal progenitors: the TN-dependent repression of dorsal progenitor identities, and the TN-independent repression of *Neurog2* and therefore neurogenesis **(Figure 2G)**. Interestingly, we found that the effects of iNkx2-2^TNmut^ most similarly recapitulated the molecular differences between human vpMNs and pMNs **(Figure 2H)**, suggesting that while Nkx2-2’s TN-dependent activity is dominant in the mouse spinal cord^16,17,22^, its TN-independent activity is dominant in human vpMNs^15,24,25^.

### TN-mutant Nkx2-2 suppresses neurogenesis without affecting dorsoventral patterning at limb-level spinal cord

While NKX2-2-expressing progenitors in the rostral cervical neural tube primarily give rise to PHOX2A^+^/ISL1^+^ branchiovisceral motor neurons, NKX2-2-expressing progenitors in more caudal levels give rise to SIM1^+^ V3 interneurons^27^ (**Figure 3A**). In order to evaluate the effects of *Nkx2-2* expression in caudal spinal progenitors, we therefore developed a new mouse ESC differentiation protocol that facilitates efficient production of brachial motor neurons.

**Figure 3:**
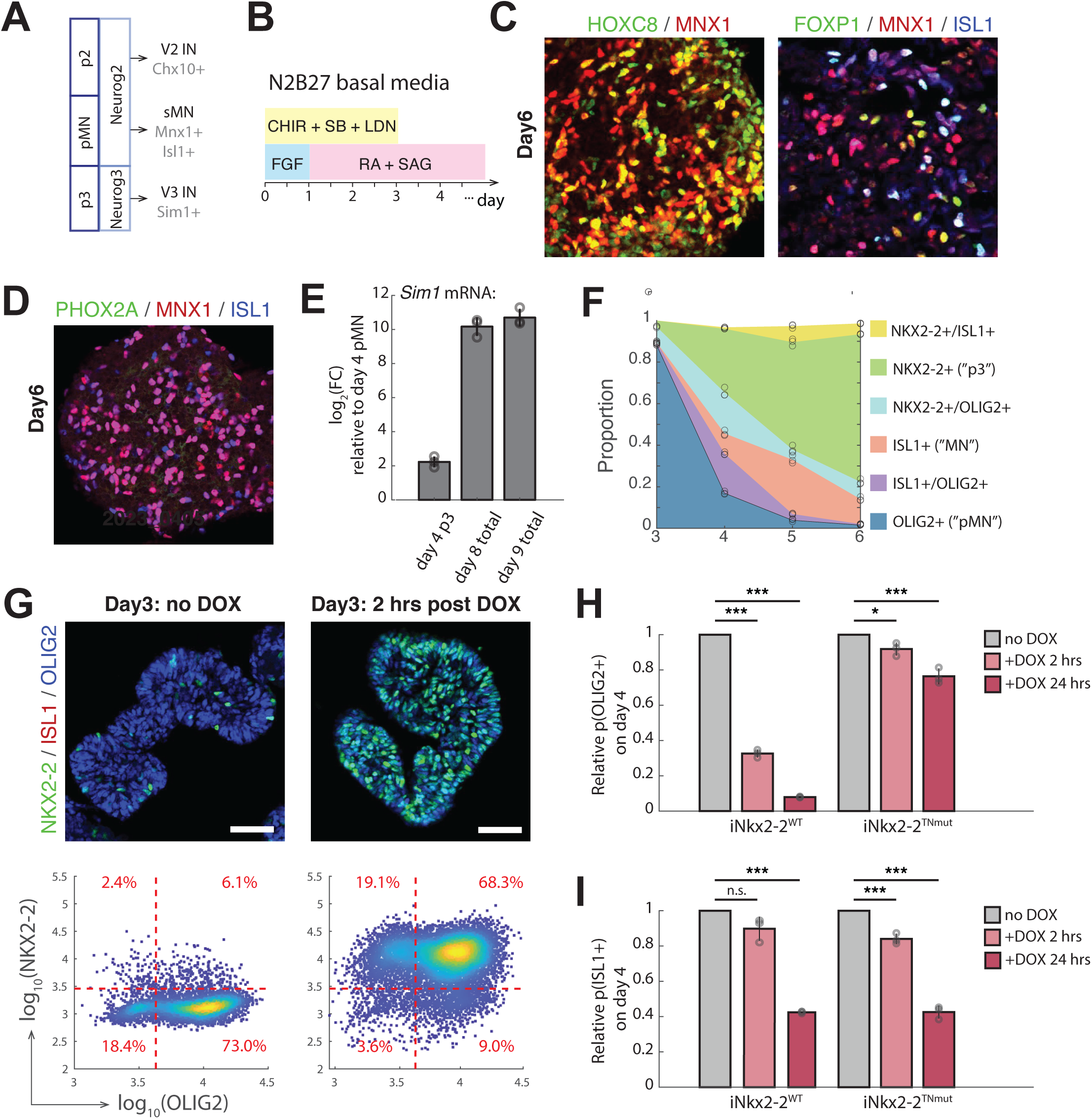
TN-mutant Nkx2-2 suppresses neurogenesis without affecting dorsoventral patterning at limb-level spinal cord. **(A)** Schematic of ventral progenitor domains in the brachial (caudal cervical) spinal cord; at this level, p3 progenitors generate V3 interneurons (IN) instead of branchiovisceral motor neurons. **(B)** Summary of culture conditions that produce brachical motor neurons. **(C)** Under these conditions, HOXC8^+^ somatic motor neurons (MNX1^+^) are produced, indicating brachial-level identity. In accordance with their brachial-level identity, a subset of these motor neurons express FOXP1, indicating that they are limb-innervating, LMC-like motor neuron subtypes. **(D)** All motor neurons (ISL1^+^) produced under these conditions are somatic (MNX1^+^). No expression of PHOX2A is observed, indicating absence of branchiovisceral motor neurons. **(E)** Relative *Sim1* mRNA levels increase dramatically over time, indicating that V3 interneurons are likely also produced at later stages under these conditions. **(F)** Summary of NKX2-2, ISL1 and OLIG2-expressing cell proportions (normalized to total number of cells expressing at least one of these three genes) over time. Day 3 shows high enrichment for pMNs. **(G)** DOX treatment on day 3 leads to rapid and uniform NKX2-2 protein expression that is acutely co-expressed with OLIG2. **(H)** OLIG2^+^ cell proportions are dramatically reduced 24 hours after DOX treatment (2 hour-pulse or sustained for 24 hours) in iNkx2-2^WT^ by day 4, but largely maintained with iNkx2-2^TNmut^. **(I)** In contrast, ISL1^+^ cell proportions are similarly reduced 24 hours after DOX treatment with both iNkx2-2^WT^ and iNkx2-2^TNmut^.

Our strategy was to approximate human pluripotent stem cell differentiation conditions that were shown to primarily give rise to HOXC8^+^ brachial motor neurons^28^. In order to best match the mouse culture conditions to our human conditions, we started with mouse pluripotent cells that were maintained in a state of “primed” pluripotency, similar to that of human pluripotent cells. We then treated the cells with dual-SMAD inhibition to promote neuronal differentiation, while activating Fgf and Wnt signaling pathways (with FGF2 and GSK3β inhibitor CHIR99021), that jointly promote the specification of a brachial neural tube identity **(Figure 3B)**. We found that the motor neurons produced under these conditions expressed HOXC8, and many acquired expression of FOXP1, a marker of limb-innervating motor neuron subtypes **(Figure 3C)**. Importantly, we did not observe any PHOX2A^+^ motor neurons under these conditions **(Figure 3D)**, but instead observed a dramatic increase in *Sim1* expression levels at later (day 8+) stages, indicative of V3 interneuron production **(Figure 3E)**.

In order to determine the dynamics and kinetics of differentiation under these new conditions, we sampled cells every 24 hours and immunolabeled them for OLIG2 (pMN marker), NKX2-2 (p3 marker), and ISL1 (motor neuron marker), assessing the proportions of cells expressing one or more of these genes via flow cytometry **(Figure 3F)**. Very few cells expressed OLIG2, NKX2-2 or ISL1 prior to day 3. OLIG2 expression peaked on day 3, and postmitotic motor neuron marker ISL1 increased progressively over the next 72 hours. p3 progenitors (NKX2-2^+^) also appeared and increased in proportion between days 3 and 6 of differentiation.

To evaluate the effects of NKX2-2 on *Olig2* expression under these conditions, we treated differentiating iNkx2-2 cell lines with DOX on day 3. We found that with either iNkx2-2^WT^ or iNkx2-2^TNmut^, NKX2-2 protein was rapidly induced in as little as 2 hours upon DOX addition in the vast majority (∼90%) of cells, creating a population that was highly enriched (68.3%) for vpMN-like (NKX2-2^+^/OLIG2^+^) cells **(Figure 3G)**. We therefore decided to remove DOX following a 2-hour (or 24-hour) pulse to see how transient Nkx2-2 induction in pMNs would affect their subsequent differentiation.

Although NKX2-2 and OLIG2 were co-expressed 2 hours after addition of DOX in both iNkx2-2^WT^ and iNkx2-2^TNmut^, co-expression was unstable in iNkx2-2^WT^, leading to downregulation of OLIG2 in most cells over the next 24 hours, whereas NKX2-2 and OLIG2 co-expression was largely maintained in iNkx2-2^TNmut^ cultures, similar to what was observed in rostral differentiation conditions **(Figure 3H)**. In contrast, ISL1^+^ motor neurons in both iNkx2-2^WT^ and iNkx2-2^TNmut^ were similarly decreased **(Figure 3I)**, confirming that under caudalized differentiation conditions as well, NKX2-2 represses motor neurogenesis in a TN-independent manner, while repressing *Olig2* and the motor neuron lineage in a TN-dependent manner.

### Nkx2-2-dependent Neurog2 repression delays and extends motor neurogenesis

We then asked whether NKX2-2 represses *Neurog2* in a TN- and Notch-independent manner at brachial levels as well. To test this directly, we treated iNkx2-2^TNmut^ cells with gamma secretase inhibitor DAPT to block Notch signaling following 3 hours of DOX induction. In non-DOX-treated controls, this led to a sharp increase in NEUROG2 levels among OLIG2^+^ progenitors. In contrast, very few cells were NEUROG2^+^ in iNkx2-2^TNmut^, confirming that NKX2-2 indeed represses NEUROG2 independently of its TN domain and the Notch signaling pathway **(Figure 4A)**.

**Figure 4:**
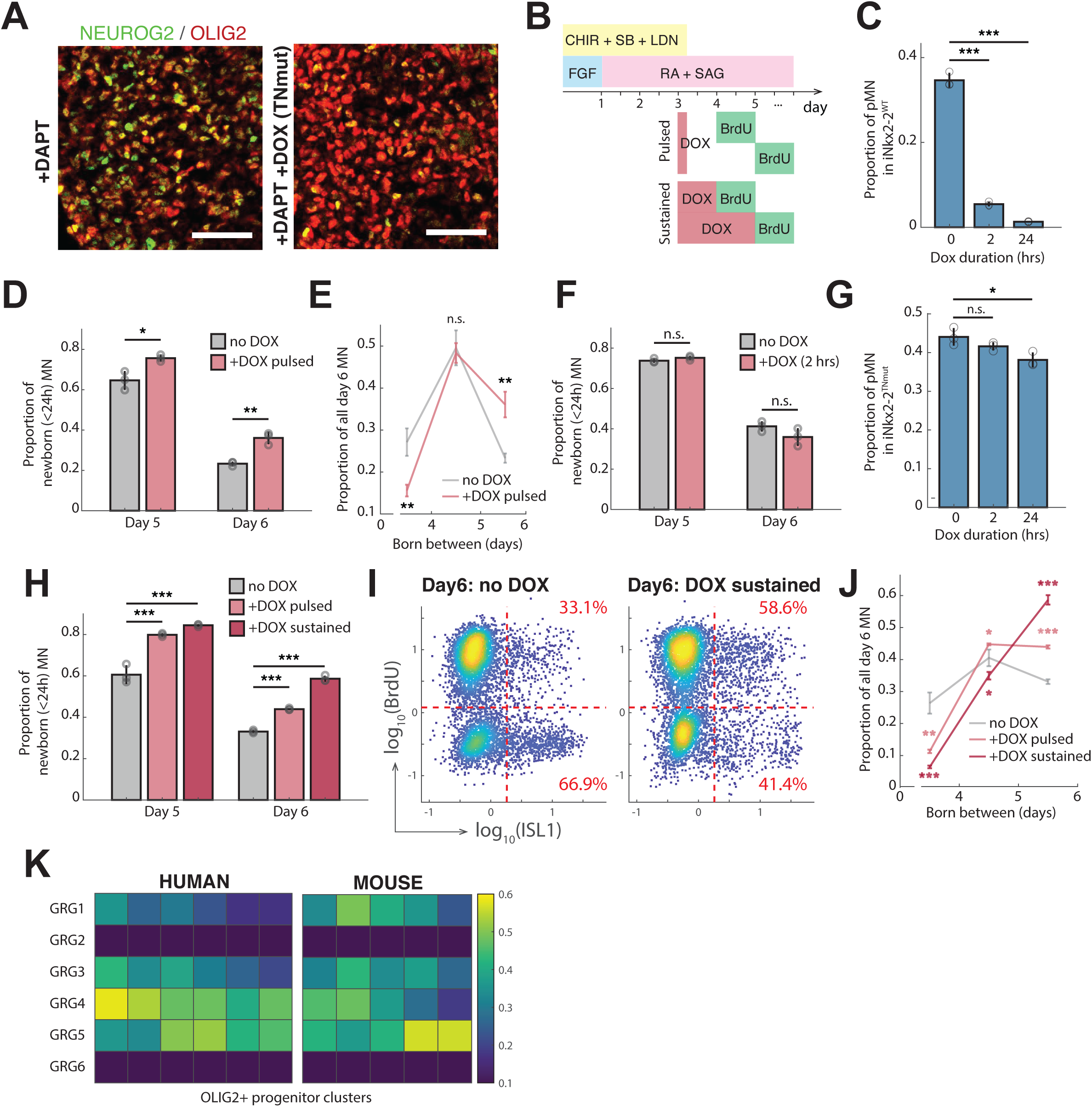
Nkx2-2-dependent Neurog2 repression delays and extends motor neurogenesis. **(A)** DAPT treatment of day 3 cultures to block Notch signaling leads to high NEUROG2 levels in OLIG2^+^ pMNs on day 4. However, iNkx2-2^TNmut^ blocks DAPT-induced NEUROG2 increase without repressing OLIG2, indicating that Nkx2-2 represses Neurog2 independently of its TN domain, Olig2, and the Notch pathway. **(B)** Summary of culture conditions for birthdating motor neurogenesis following DOX induction of *Nkx2-2*. **(C)** Ectopic expression of WT *Nkx2-2* leads to dose-dependent loss of pMNs. **(D)** DOX pulse in iNkx2-2^WT^ leads to slightly but significantly delayed motor neurogenesis. **(E)** DOX pulse in iNkx2-2^WT^ reduces proportion of earlier-born (before day 4) motor neurons, and increases proportion of later-born (after day 5) motor neurons. **(F)** DOX pulse in parental mouse ES cell line with no DOX-inducible *Nkx2-2* has no effect on the timing of motor neurogenesis. **(G)** Ectopic expression of TN-mutant *Nkx2-2* largely maintains pMN proportions. **(H)** DOX induction in iNkx2-2^TNmut^ delays motor neurogenesis in a dose-dependent manner. **(I)** Among day 6 motor neurons (ISL1^+^), the majority of them are born prior to day 5 (BrdU-negative). However, with sustained ectopic expression of TN-mutant *Nkx2-2*, almost 60% of motor neurons are born after day 5 (BrdU^+^). **(J)** Sustained ectopic expression of TN-mutant *Nkx2-2* not only delays but also extends motor neurogenesis, as evidenced by its monotonically increasing birthcurve, in contrast to the control birthcurve, which peaks between days 4 and 5 and decreases thereafter. **(K)** Normalized expression levels of Groucho genes in human and mouse *Olig2*-expressing progenitor clusters derived from single-cell RNA-sequencing of in vitro motor neurogenesis^15^, sorted from left to right with increasing *Nkx2-2* expression levels.

Finally, we tested whether *Neurog2* repression by ectopic *Nkx2-2* expression could functionally mimic vpMN neurogenic behavior by delaying and extending neurogenesis in mouse pMNs. We first tested this in iNkx2-2^WT^. To circumvent complete loss of OLIG2 expression, we pulsed cultures with 2 hours of DOX on day 3, and added BrdU on day 4 or 5, prior to fixing cells on day 5 or 6 to label motor neurons born within the last 24 hours **(Figure 4B)**. Although DOX pulse led to a substantial decrease in motor neuron progenitors **(Figure 4C)**, iNkx2-2^WT^ had a small but significant effect on the birth times of motor neurons that were specified despite Nkx2-2 induction **(Figure 4D)**. We saw that with a 2-hour DOX pulse in iNkx2-2^WT^, the proportion of earlier-born (prior to day 4) motor neurons decreased, and that of later-born (after day 5) motor neurons increased **(Figure 4E)**. To test that this effect was not caused by DOX per se, we performed the same experiment with the parental mouse cell line lacking the ectopic *Nkx2-2* cassette, and saw that DOX pulse did not affect motor neuron birth times in the absence of *Nkx2-2* induction **(Figure 4F)**.

Next, we tested whether we could further extend and delay motor neurogenesis using iNkx2-2^TNmut^. Because TN-mutant NKX2-2 does not suppress the motor neuron lineage but still represses *Neurog2*, we reasoned that with iNkx2-2^TNmut^, we could apply sustained DOX induction and therefore more effectively suppress neurogenesis while circumventing loss of OLIG2^+^ progenitors. Indeed, we found that in iNkx2-2^TNmut^, the vast majority (>90%) of pMNs remained OLIG2^+^, even following 24-hour DOX induction **(Figure 4G)**. Therefore, in addition to performing the same 2-hour DOX pulse experiment, we also treated cultures to sustained DOX treatment, in which we added DOX on day 3 for: (a) 24 hours and assessed its effect on motor neurons born between days 4-5, or (b) 48 hours and assessed its effect on motor neurons born between days 5-6 **(Figure 4B)**.

Similar to what we had observed with iNkx2-2^WT^, we found that motor neurogenesis was significantly delayed already with a 2-hour DOX pulse in iNkx2-2^TNmut^ **(Figure 4H)**. Interestingly, iNkx2-2^TNmut^ had an even greater effect on motor neuron birth times than iNkx2-2^WT^, especially on earlier neurogenesis (days 4-5). This may be because cells that failed to induce Nkx2-2 (and therefore undergo earlier neurogenesis) are likely over-represented in the motor neuron population in iNkx2-2^WT^. When we calculated the birth curves for iNkx2-2^TNmut^ with DOX pulse, we saw that early (before day 4) motor neurogenesis was decreased and late (after day 5) motor neurogenesis was increased, similar to the effect seen with DOX pulse in iNkx2-2^WT^ **(Figure 4J)**.

Cultures treated with sustained DOX exhibited an even greater effect on motor neurogenesis. Sustained, 48-hour DOX more than doubled the effect on later (day 5-6) neurogenesis compared to DOX pulse (33% increase with DOX pulse, 77% increase with sustained DOX), **(Figure 4H)**. Whereas only a third of day 6 motor neurons were born after day 5 in the no-DOX control, nearly 60% of cells were born after day 6 with 48 hours of DOX induction **(Figure 4I)**. Remarkably, sustained DOX induction in iNkx2-2^TNmut^ produced a birthcurve in which birth rates monotonically increased over time, whereas in all other conditions, the birthcurve peaked between days 4-5 and decreased thereafter **(Figure 4J)**. The dramatically different shape of the birthcurve following sustained DOX induction further indicated that neurogenesis is not only delayed, but also extended in iNkx2-2^TNmut^, similar to vpMNs. Together, these results confirm that Nkx2-2 extends motor neurogenesis in a TN-independent manner, recapitulating the behavior of human vpMNs in mouse pMNs.

## DISCUSSION

The mechanisms that lead to neural expansion in humans and primates has been the subject of intense investigation. Recent studies underscore the diversity of mechanisms that evolution has produced and modified that lead to extended and expanded neurogenesis in the human CNS ^4,6,7,13,15,29,30^. Our studies reveal a surprising and distinguishing aspect of the spinal motor neuron lineage, which is that its evolutionary extension of neurogenesis in humans arises through changes in the interaction between two genes, *NKX2-2* and *OLIG2*. While NKX2-2 strongly and rapidly represses *Olig2* in rodents and chicks through its repressive TN domain during neurogenesis^22^, this repressive activity seems to be diminished or absent in the early human spinal cord^15,24,25^. This difference in turn allows for co-expression of *NKX2-2* and *NEUROG2* in the human spinal motor neuron lineage, thus revealing NKX2-2’s TN-independent repression of *NEUROG2* that is masked in rodents and chicks.

While the mitotic behavior of progenitors throughout the spinal cord are affected by the Notch signaling pathway, motor neuron and p3 progenitors utilize different Neurog paralogs in their Hes cycle: motor neuron progenitors (i.e., both pMNs and vpMNs) express Neurog2, and p3s express Neurog3. In pMNs, *Olig2* has been shown to repress Hes1/5, thereby shifting the Hes cycle balance toward increased Neurog2 and neurogenesis ^17,31–33^. Because of this, motor neurons are some of the earliest-born neurons in the CNS^34^. Our studies show that NKX2-2 can also repress *Neurog2* independently of *Olig2* and the Notch pathway. While this effect is normally not observed in rodents and chicks, it is apparent in humans and unmasked with TN-mutant *Nkx2-2* in mouse systems. What mechanism underlies this TN-independent repression remains unknown, but it is interesting to consider whether the NK2-specific domain (SD) might be required for this activity, as this domain has been shown to play an important role during cell fate specification in the developing pancreas^20^. Testing this will require mutating both the SD and TN domain, as TN-dependent *Olig2* repression will compensate for the potential loss of *Neurog2* repression in the SD mutant.

In light of these observations, one outstanding question is why NKX2-2’s TN-dependent repression of *OLIG2* appears to be diminished or absent in humans. Both human and mouse orthologs of *NKX2-2* harbor the same TN domain, and only 3 of 273 amino acid residues differ across the rest of the protein^35^. Although previous studies have shown that NKX2-2 interacts with the Groucho family of co-repressors through its TN domain to suppress the motor neuron lineage^22^, both human and mouse motor neuron progenitors have similar expression patterns of Groucho genes **(Figure 4K)**, raising the question of whether species-specific differences in posttranslational modifications or levels of other NKX2-2 interactors attenuate the TN-dependent activity in the human neural tube.

In mice and chicks, *Nkx2-2* and *Olig2* co-expression occurs later in oligodendrocyte precursor cells (OPCs) that arise from the ventral spinal cord, following motor neurogenesis. It was previously shown that while ectopic expression of *Nkx2-2* represses *Olig2* and converts pMNs into p3 progenitors^16,17,22^, ectopic expression of both *Nkx2-2* and *Olig2* leads to premature expression of OPC marker *Sox10* in early chick pMNs^33^. Our studies show that even though stable co-expression of *Nkx2-2* and *Olig2* is achieved with TN-mutant *Nkx2-2*, it does not seem to divert progenitors into a gliogenic fate, as they still give rise to motor neurons, albeit in a delayed and protracted manner, and Sox10 expression is not increased. Nevertheless, it remains to be determined whether manipulations that lead to premature and stable co-expression of *Nkx2-2* and *Olig2* in mouse pMNs affect the specification of OPCs later.

Our studies indicate that NKX2-2 represses *NEUROG2* expression and extends motor neurogenesis. Previously, we showed that due to their delayed and extended neurogenic timeline, vpMNs give rise to ∼10 motor neurons per progenitor while pMNs give rise to ∼2^15^. When NKX2-2 is knocked out, or in mouse systems where vpMNs don’t exist, the motor neuron lineage produces ∼2 motor neurons per progenitor, indicating that NKX2-2 is the distinguishing factor in the functional difference between human and mouse motor neuron progenitors. Although we were eager to test whether ectopic expression of *Nkx2-2* would lead to increased motor neuron production in mouse cultures, we found that ectopic expression of *Nkx2-2* (both TN-mutant and WT) led to slightly smaller embryoid bodies. This effect was further exacerbated with sustained and late DOX treatment, and especially when combined with DAPT treatment, suggesting that ectopic expression of *Nkx2-2* may be toxic for motor neurons, in particular. Given these limitations, it will be important to devise new strategies that can lead to progenitor-specific expression of TN-mutant *Nkx2-2*, to see if we can then observe vpMN-like cells that not only extend but also expand motor neurogenesis in mouse.

## AUTHOR CONTRIBUTIONS

Conceptualization: SJ, HW

Experiments: SJ, EA, JD

Data analysis: SJ

Writing – original draft: SJ

Writing – review & editing: SJ, HW

## ACKNOWLEDGEMENT

We would like to thank Dr. Lori Sussel for feedback and discussion, Dr. Michael Closser and Dr. David Gifford for help with and access to shared computing. We would also like to thank Michael Kissner for assistance with flow cytometry.

## Funding

Jerry and Emily Spiegel endowed chair (HW)

National Institutes of Health grant R01NS116141 (HW)

National Institutes of Health grant R01NS089676 (HW)

Project ALS (HW)

Columbia Stem Cell Initiative seed grant (SJ)

National Institutes of Health grant K99MH130892 (SJ)

## MATERIAL AND METHODS

### Pluripotent stem cell culture

Human iPS cells (NCRM-1 background: National Institute of Health - Center for Regenerative Medicine) were grown on Matrigel (Corning CLS354277)-coated plates in mTeSR-Plus (STEMCELL Technologies 100-0276) or on a bed of irradiated mouse embryonic fibroblasts (Thermo Fisher A34180), and differentiated according to previously established protocols^28^, without DAPT.

Mouse ES cells^36^ were grown on gelatin-coated plates in LIF+2i-containing basal media^37^. Mouse cells were differentiated toward rostral cervical motor neurons as previously described^26^. For caudal differentiation, mouse ES cells were first differentiated toward (and maintained in) primed pluripotency following previously described conditions^38^. To initiate differentiation, primed pluripotent mouse cells were dissociated, seeded at 500K cells per 6-cm plate in N2 (Thermo Fisher) and B27 (Thermo Fisher) containing media with 10ng/mL Fgf2, 10uM Y27632, 3uM CHIR99021, 20uM SB431542, 0.1uM LDN193189. 24 hours later, on day 1, cells were transferred to a 10-cm plate with N2B27 media with 0.1uM retinoid acid, 0.1uM Smoothened agonist (SAG), 3uM CHIR99021, 20uM SB431542, 0.1uM LND193189. On day 3, media was changed to N2B27 with 0.1uM retinoid acid and 0.1uM SAG.

10uM BrdU was added to cultures for birth-dating. DAPT was used at 10uM to block Notch signaling. 1ug/mL doxycycline was added to induce ectopic Nkx2-2 expression.

All cell culture experiments were repeated across at least three biological replicates (each consisting of independent embryoid bodies derived from different wells or plates) unless otherwise stated.

#### Generation of inducible Nkx2.2 cell lines

Inducible ESC lines were generated as previously described^39^. Briefly, open reading frames of genes were cloned by PCR from pcDNA3-WT Nkx2.2^40^ or pcDNA3-TN mutant Nkx2-2^41^. To minimize the introduction of mutations during PCR amplification, Phusion polymerase was used (New England Biolabs, M0530S). Open reading frames were directionally inserted into pENTR/D-TOPO vector (Invitrogen, K2435-20) following the manufacturer’s instructions. The 5′ primer always contained the added CACC sequence to ensure directional integration, and the 3’ primer included the stop codon. To generate Flag-tagged protein sequences, the pENTR plasmid with stop codon was recombined with p2Lox-FlagB plasmid^39^ using Gateway LR Clonase II enzyme mix (Invitrogen, 11791-020) per the manufacturer’s instructions. Inducible lines were generated by treating the recipient 2lox.Cre ESCs^36,39^ for 16 h with DOX (1μg/mL) to induce Cre recombinase expression followed by nucleofection (Lonza, VPG-1004) of the p2Lox-FlagB plasmids containing the desired construct. After selection with G418 (Sigma-Aldrich, A1720), resistant clones were pooled, characterized, and expanded. New inducible lines generated in this study include those harboring the wildtype *Nkx2-2* (iNkx2.2^WT^) and the TN mutant *Nk2-2* (iNkx2-2^TNmut^) transgenes.

#### RNA-seq sample preparation and alignment

Human data: day 11 cultures were dissociated, fixed with 4% PFA for 15 minutes, after which they were permeabilized with ice-cold methanol and rehydrated with 1% BSA in PBS. Cells were then immunostained for NKX2-2, OLIG2 and ISL1/2 and FAC-sorted for NKX2-2^+^/OLIG2^+^/ISL1/2^−^ (vpMN) and NKX2-2-/OLIG2^+^/ISL1/2^−^ (pMN) populations. RNA was then purified from these sorted populations following previously described protocols^42^.

Mouse data: RNA was extracted from day4 embryoid bodies using Trizol Reagent (Life Technologies, SC: 15596-018) in combination with RNeasy Mini Kit (Qiagen, 74104). Libraries were prepared with Illumina TrueSeq RNA Prep Kit. Inclusion of samples required RNA integrity (RIN) values > 8.0 as determined with Agilent Bioanalyzer 2100. RNA was sequenced using Illumina HiSeq2000 at the Columbia Genome Center, and three replicates were obtained per condition. Reads were mapped to the mouse genome (mm10) using the RNA-seq alignment algorithm in the STAR (Spliced Transcripts Alignment to a Reference) software package (version 2.3.1)^43^.

#### RNA-seq analysis

Differential RNA expression across cohorts was assessed using EdgeR^44^ (version 3.30.3). For human data, raw counts from two biological replicates of fixed, stained, and FACS-purified day 11 vpMN and pMNs were used. For mouse data, raw counts from replicate samples in the non-dox-treated control, iNkx2.2^WT^, iNkx2-2^SDmut^, and iNkx2-2^TNmut^ groups were analyzed.

#### Calculation of birthcurve

Cells were treated with BrdU on day 4 and collected on day 5, or treated with BrdU on day 5 and collected on day 6. The proportions of BrdU-positive MNX1^+^ motor neurons, *p*(*BrdU*^+^|*MN*_*day* 5_)) and *p*(*BrdU*^+^|*MN*_*day* 6_), were calculated by flow cytometry following immunofluorescence staining and BrdU-labeling. From this, we calculated the birthcurve for motor neurons present at day 6:

Born before day 4: (1 − *p*(*BrdU*^+^|*MN*_*day*6_)) × (1 − *p*(*BrdU*^+^|*MN*_*day*5_))

Born between days 4 and 5: (1 − *p*(*BrdU*^+^|*MN*_*day*6_)) × *p*(*BrdU*^+^|*MN*_*day*5_)

Born after day 5: (*BrdU*^+^|*MN*_*day*6_)

#### Immunohistochemistry

##### For flow cytometry or FACS

Embryoid bodies (at least 50 EBs per sample) were pooled and dissociated using papain following the Neural Tissue Dissociation Kit – P (Miltenyi Biotec 130-092-628) and quenched with ovomucoid protease inhibitor (Worthington LK003182). Following dissociation, cells were washed twice with 1X PBS, fixed using 4% paraformaldehyde, washed with PBS, and then permeabilized with ice-cold methanol. Permeabilized cells were rehydrated with 1% BSA in PBS solution, and subsequently immunostained using a standard two-step immunofluorescence labeling method.

##### Sectioned embryoid bodies

Whole embryoid bodies were fixed in 4% paraformaldehyde, washed several times with PBS, then put through 30% sucrose prior to embedding in OCT and freezing. Frozen blocks were sectioned in a cryostat to 15um slices and mounted on microscope slides (Superfrost Plus, Fisher 12-550-15). Sections were rehydrated with staining solution (1% BSA, 0.1% Triton X-100 in PBS), then immunostained using a standard two-step immunofluorescence labeling method.

##### Primary Antibodies; host species and concentrations used

ISL1 (Goat; 1:5000; Neuromics Cat# GT15051-100, RRID:AB_2126323; has 10% cross-reactivity to ISL2), MNX1 (Guinea pig; 1:100; from Jessell Lab), FOXP1 (Mouse; 1:400; Santa Cruz Cat# sc-398811), NKX2-2 (Mouse; 1:100; DSHB Cat# 74.5A5, RRID:AB_531794), BrdU (Rat; 1:400; Abcam Cat# ab6326, RRID:AB_305426), OLIG2 (Guinea pig; 1:100; from Jessell Lab), Cleaved Caspase3 (Rabbit; 1:200; Cell Signaling #9661).

